# *In-silico* and *in-vitro* morphometric analysis of intestinal organoids

**DOI:** 10.1101/2022.12.08.519603

**Authors:** Sandra Montes-Olivas, Danny Legge, Abbie Lund, Alexander G. Fletcher, Ann C. Williams, Lucia Marucci, Martin Homer

## Abstract

Organoids offer a powerful model to study cellular self-organisation, the growth of specific tissue morphologies *in-vitro*, and to assess potential medical therapies. However, the intrinsic mechanisms of these systems are not entirely understood yet, which can result in variability of organoids due to differences in culture conditions and basement membrane extracts used. Improving the standardisation of organoid cultures is essential for their implementation in clinical protocols. Developing tools to assess and predict the behaviour of these systems may produce a more robust and standardised biological model to perform accurate clinical studies. Here we developed an algorithm to automate crypt-like structure counting on intestinal organoids in both *in-vitro* and *in-silico* images. In addition, we modified an existing two-dimensional agent-based mathematical model of intestinal organoids to better describe the system physiology, and evaluated its ability to replicate budding structures compared to new experimental data we generated. The crypt-counting algorithm proved useful in approximating the average budding structures found in our *in-vitro* intestinal organoid culture images on days 3 and 7 after seeding. Our changes to the *in-silico* model maintain the potential to produce simulations that replicate the number of budding structures found on days 5 and 7 of *in-vitro* data. The present study aims to aid in quantifying key morphological structures and provide a method to compare both *in-vitro* and *in-silico* experiments. Our results could be extended later to 3D *in-silico* models.

## 1. Introduction

Organoids are three-dimensional cell cultures capable of self-organisation which can replicate organ-specific shapes, such as crypt-like invaginations in the case of intestinal organoids. Organoids present a powerful tool for studying morphological phenotypes in diverse tissues such as the brain [1–3], intestine [4–6], and kidney [7, 8], among others. However, the mechanisms underlying the generation of heterogeneous morphologies in organoids are not entirely understood, and thus, it is still a challenge to standardise their cultures for experimental manipulation. The development of mathematical models capable of simulating the spatiotemporal dynamics of these complex systems may help to clarify the mechanisms that give rise to the tissue-specific organoid morphology. Mathematical models could also aid in standardising culture protocols, increase experimental efficiency, and lead to improved clinical treatments and regenerative therapies [9, 10]. Nonetheless, mathematical models require quantitative data to be calibrated and tested.

Due to their self-organisational and heterogeneous properties, the morphological measurement and analysis of organoids can be demanding, particularly if no staining or fluorescence is added to the culture and only bright-field images are obtained. The complexity of the 3D environment within an organoid provides a more faithful recapitulation of *in-vivo* conditions than other *in-vitro* approaches. However, the 3D structure can make image analysis of organoids challenging.

To date, organoid morphology has been quantified by segmenting *in-vitro* images and extracting basic morphometric measurements. Segmentation is a fundamental step in the extraction and analysis of morphological information. Several segmentation algorithms can be implemented on microscopy images, depending on the image quality and other properties. Among those available are region growing, seeded watershed, k-means clustering, and active contours [11]. Several segmentation methods have been developed for labelled and label-free 2D and 3D cultures [6, 12–15]. Once an image is segmented, typical morphology metrics such as diameter, perimeter, length, area, circularity, curvature and sphericity can be easily calculated [16]. However, there is still a need for algorithms that are capable of aiding in the automatic detection and quantification of prominent tissue-specific features, such as crypt-like domains in intestinal organoids.

The intestinal organoid culture, described by Sato et al. [17], was the first three-dimensional cell culture model capable of self-organising and producing intestinal crypt- and villus-like compartments. The crypt-like units contain the stem-cell niche, while most differentiated cells reside in the villus-like sections. The crypt-villus structures are the most critical characteristics of the intestinal epithelial morphogenesis [18]. The presence and frequency of crypt budding is a crucial parameter in the evaluation of organoid-forming efficiency [19]. Similarly, this parameter can be used to determine the maturity of an organoid during experimentation [20]. Usually, for these cultures, the experimentalist manually quantifies the number of crypts per organoid by observing a small set of organoids or measuring circularity as a proxy to infer crypt growth.

In recent years, there have been a great development of mathematical and computational models of organoid growth [21]. Several models have been developed to understand relations between signalling pathways, cell differentiation regulators, and their effect in the growth and shapes of intestinal organoids using different simulation frameworks [22–24]. However, not many models have focused on possible biomechanical cell interactions involved in the creation of crypt-like structures. Langlands et. al. [25] proposed a model in which crypt fission results from the interaction of two different cell populations with different stiffnesses, which the authors related to stem cells and Paneth cells. This model was developed using the Chaste framework [26, 27], which is a popular software library that can be employed to simulate multicellular tissue populations [28, 29]. Model simulations were compared to *in-vitro* experiments by measuring the organoid’s circularity. Almet et. al. [30] further explored the biomechanical properties involved in crypt fission in this model by performing a parameter sweep over the hard cells, stiffness ratio and their target proportion. They found that these two parameters affect the deformation of the epithelial monolayer and the generation of crypt-like structures. However, in both studies, the model’s assumptions for cell proliferation were not based on an specific cell type characterisation. Similarly to the previous study, they employed circularity to determine crypt fission occurrence in their simulations.

Nevertheless, relying on the circularity measure to indirectly quantify the number of crypts does not accurately quantify organoid growth once they lose their initial spheroid shape. To address this, here we present an algorithm that uses both segmented *in-vitro* and *in-silico* organoid images and retrieves an approximated number of crypts per organoid. Additionally, we propose modifications to the model proposed by Langlands et al. [25] to improve its biological basis and we compare both original and improved model simulations to *in-vitro* organoid images. Our results should provide insights into the multiscale processes orchestrating intestinal organoids growth.

## 2. Methods

### (a) Organoid culture

Intestinal cells were isolated from six week old C57BL/6 mice and cultured as previously described (adapted from the method of Sato & Clevers [31]). In brief, 10cm of proximal small intestine was removed from mice and villi removed by scraping. The remaining intestine was washed 5x in cold PBS before incubation in PBS containing 2mM EDTA. Crypts were mechanically dissociated, and crypt fractions isolated by centrifugation at 600 rpm for three minutes before being mixed with Matrigel and pipetted into prewarmed 24 well plates, 50 *µ*L Matrigel per well. Plates were incubated at 37°C for 10 minutes for Matrigel to polymerize after which Advanced DMEM:F12 (ADF; Gibco) media was added (500 *µ*L per well) [supplemented with 0.1% BSA (Sigma-Aldrich), 2mM glutamine (Gibco), 10mM HEPES (Sigma-Aldrich), 100 units/ml penicillin, 100 units/ml streptomycin, 1% N2 (Gibco), 2% B27 (Gibco) and 0.2% N-acetyl-cysteine (Gibco)] with the addition of 50ng/ml EGF (Peprotech), 100 ng/ml Noggin (Peprotech, London, UK) and 500ng/ml mR-Spondin 3 (R&D systems, MN, USA). Crypts were then cultured for 2-3 days before organoids were formed. Medium was changed every 4 days.

### (b) Segmentation

Brightfield images were acquired on days 3, 5 and 7 of intestinal organoid culture, maintaining the samples at 37°C, using a Leica DMI6000 inverted microscope with 5x and 10x magnification lenses from the Wolfson Bioimaging Facility, University of Bristol. Images were acquired as 30 z stacks to cover the depth (z-axis) of the Matrigel™ domes. Figure 1 presents sample images of the organoid cultures obtained on experimental days 3, 5, and 7 after seeding.

**Figure 1:**
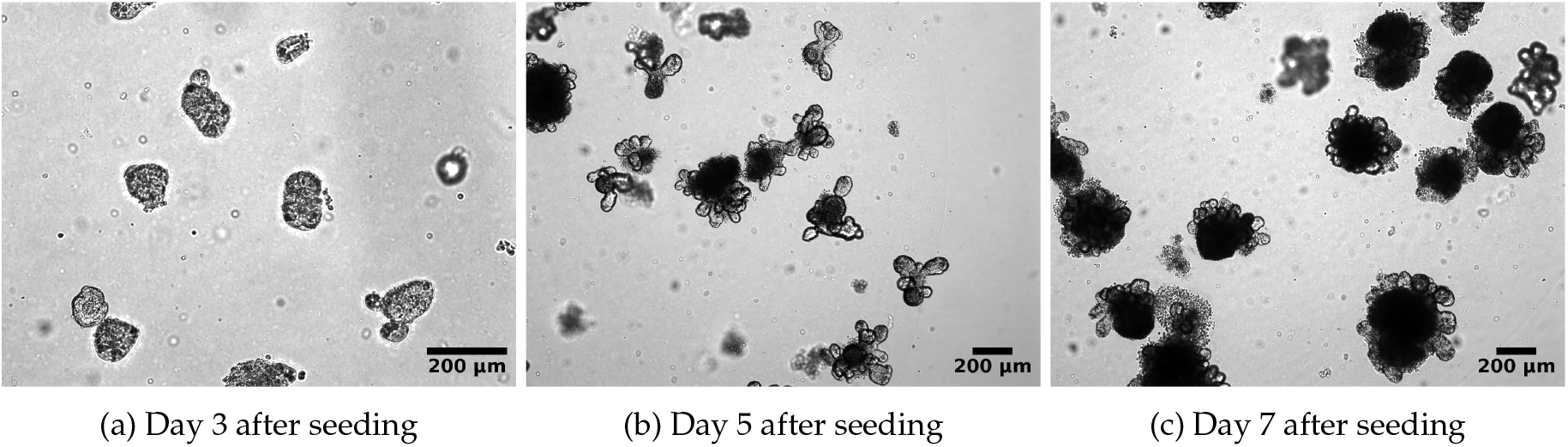
Timeline images of *in-vitro* intestinal organoid culture. Samples of *in-vitro* intestinal organoid culture stacked images obtained at days 3 (a), 5 (b) and 7 (c) after seeding. 10x (a) and 5x (b and c) objective. Scale bar: 200 *µ*m.

First, all images obtained from the microscope were pre-processed by stacking the focal planes in the z-plane to get a single focus image using a custom code. Second, two segmentation methods were compared to assess the effect of segmentation quality on the proposed automatic crypt quantification: (i) manual free-hand segmentation and (ii) an open-source segmentation software specially developed for analysing organoid cultures.

#### (i) Manual segmentation

The manual segmentation was performed in MATLAB using the in-built function *imageSegmenter* and the assisted freehand to select the regions of interest (ROI). Viable organoids were selected through visual inspection by the user.

#### (ii) Open source-segmentation software for organoids

OrganoSeg is a software developed using the Image Processing Toolbox from MATLAB [32]. To use this software, the images (grouped by date) were used as input and the values of intensity threshold (0.1–0.7), max window size (100–500 pixels) and size-exclusion threshold (500–2500 pixels) were selected for each image separately.

The OrganoSeg software presents a few advantages compared to the previous method (i) as it was developed especially for organoid segmentation. One of these advantages is the option to detect all elements that comply with the selected parameters. The user is then given the option to remove detected objects that the user can visually classify as wrong. Another useful option that the interface includes is the ability to select an out-of-focus correction, which can be especially useful for focal plane stacked images. However, using this method makes it difficult to segment organoids that overlap or touch each other.

All output masks from both methods were saved for later analysis. The output masks contained those segmented organoids found to be viable. The user classified whether each organoid was viable by visually considering if the membrane on the organoid appeared intact and there were no breakage and leak of cells into the Matrigel™.

### (c) *In-silico* model

#### (i) Langlands et al. model

The first computational model that explored the effect of heterogeneous biomechanical cell properties on the development of crypts in intestinal organoids was presented in a study performed by Langlands et al. [25]. This model was later extended by Almet et al. [30] to explore further biomechanical properties involved in crypt fission. The general premise of these 2D cell-based models was to test if crypt fission could be initiated solely by different stiffness of two cell types, one soft and another hard, rather than by a signalling cue.

The model in [25] represents a confluent epithelial monolayer (i.e. a cross-section of a 3D intestinal organoid) and was developed using the agent-based computational framework Chaste [26, 33]. The monolayer is defined by a set of epithelial cells delineated by a honeycomb mesh created using Voronoi Tessellation and Delaunay triangulation. In the system, cells become neighbours and connected if they share an edge in the Delaunay triangulation. A cell can lose contact with another one if a set cut-off length distance is exceeded. The epithelial layer is formed by two cell types, which differ only in their relative stiffness. During the simulation, cells are exposed to interactive forces between their neighbours and the simulated Matrigel™ through a linear spring force. For more information about parameter values please see Table 1.

**Table 1:** Models parameters and values. The majority of parameter values are maintained between the original Langlands et al. and our modified unless stated otherwise. Distances are measured by cell diameters (CD), and time is scaled in hours.

| Parameter | Value |  | Units | Reference |
| --- | --- | --- | --- | --- |
|  | Langlands et al. | Proposed model |  |  |
| Mature cell spring rest length | 1 | Maintained | Cell diameter (CD) | [28, 30, 70] |
| Spring stiffness | 15 | Maintained | $N \text{ CD}^{-1}$ | [28] |
| Constant drag coefficient | 1 | Maintained | $N \text{ hours CD}^{-1}$ | [30, 71, 72] |
| Stiffness ratio of TAs | - | 4.5 | Relative to spring stiffness | Assumed |
| Stiffness ratio of PCs | 4.5 | Maintained | Relative to spring stiffness | [25, 30] |
| Basement membrane stiffness | 10 | Maintained | $N \text{ CD}^{-1}$ | [30] |
| Target curvature | 0.2 | Maintained | $\text{CD}^{-1}$ | [30] |
| Initial length of epithelial monolayer | 20 | Maintained | In CD | - |
| G1 phase of cell-cycle length | U(12, 14); all cells | U(12, 14); SCs and TAs only | In hours | [30] |
| Differentiated cells cell-cycle length | - | Inf | Do not divide | [35, 36] |
| Total simulation time | 168 | Maintained | In hours | - |
| Timestep | 0.005 | Maintained | In hours | [28, 30] |

The proliferation of all epithelial cells in this model is defined by a stochastic cell cycle duration sampled from a uniform distribution of U(12,14) hours. When a cell divides, two daughter cells connected by a spring laying in a randomly chosen direction instantly replace the original cell. The daughter cells are initially positioned in opposing orientations, 0.05 cell diameters distant from the original parent cell, and the new spring connection increases linearly from 0.1 to 1 during the first hour of cell-cycle. A user-defined division parameter controls the probability of soft cell self-renewal in the model. All epithelial cells generate one daughter cell that is identical to the parent and a new cell with probability *p* of being a soft cell and probability *p-1* of being a hard cell (see Figure 4a). Thus, both cell types will always be present in the system according to the set proportion. Additionally, all epithelial cells divide *ad infinitum*. Cell death is defined by anoikis: any epithelial cell that loses contact with its neighbours and enters the lumen or the Matrigel™ is removed from the simulation. Otherwise, if the epithelial cell is not separated from the layer, it can keep dividing according to the set proportion.

This model has proved capable of mimicking crypt formation, showcasing the effect of biomechanical properties, like stiffness and the cellular target proportions of soft and hard cells required to promote crypt fission using only two cell populations [25, 30]. However, there is still room for improvement, as its assumptions for cell proliferation are not based on a specific cell type characterisation. The first study that describes the model [25] refers to another model by Pin et al. [34], in which the Young modulus (stiffness) of Paneth cells was approximated in comparison to that of stem cells. Their results showed that Paneth cells are approximately four times stiffer than stem cells. Thus, it could be assumed that stem and Paneth cells are soft and hard cells in the model, respectively. However, the model does not take into account the presence of other intestinal cells such as enteroendocrine (EEC), enterocyte (EC), transit-amplifying cells (TA), goblet (GC) and Tuft (TC) cells. Additionally, it is known that Paneth cells are terminally differentiated cells and not capable of producing stem cells nor other Paneth cells [35, 36]. Therefore, with the current 2D model configuration it would be challenging to produce crypt fission if Paneth cells were the most stiff cells in the system, as other cells would not be capable to push or move them.

Therefore, we propose a few modifications that aim to enhance the model’s representation of biological basis such as the initial cell population proportions and the model’s cell proliferation. These modifications are further explained in section 3 (a).

### (d) Crypt counting method

The organoids’ masks must be processed to obtain possible crypt sections. Figures S1 and S2 summarise the crypt counting method processes applied to *in-vitro* and *in-silico* organoid images respectively. One important difference between the *in-vitro* and *in-silico* data is the method of extraction used. While *in-vitro* organoid images are segmented to obtain the organoid’s boundary (see Figure S1b) and the boundary of each organoid was extracted using *bwboundaries* function of MATLAB (see Figure S1c), *in-silico* organoids’ boundaries are obtained by collecting the cells’ centre points (see Figure S2b), and applying a genetic algorithm of the travelling salesman problem [37] to obtain the organoid’s boundary (see Figure S2c).

Each raw boundary is estimated using a Fourier approximation [38] to soften the mask’s rough edges acquired during the segmentation image processing (see Figure S1d and Figure S2d). It is expected that the required number of harmonics used to approximate the boundaries would change between experimental days as organoids grow and have more crypts with less definition reflected in the mask. In the images obtained from three day old organoids, it can be observed that most organoids have between 1 and 2 crypts, and the arc length of the crypts is relatively large compared to the main body of the organoid. Therefore, these images do not require high precision, and a lower number of harmonics suffice, as it produces a softer boundary approximation. However, as organoids grow and produce more crypts, these cannot be defined as clearly in the obtained masks, as can be observed in Figure 1 of our experimental data for days 5 and 7. In these cases, small changes in curvatures can indicate the presence of a crypt, and the Fourier approximation is required to be more precise. Thus, images obtained on days 5 and 7 of the experiment would need a higher number of harmonics. Therefore, such number was included as a main parameters of the algorithm used to detect crypts and required optimisation.

In the present study, we used the approximated boundary and calculated its curvature and normal. The curvature and normal values were then used to modify the boundary and obtain all convex and concave regions (see Figures S1e and S2e). A middle point was calculated for all concave regions (inward sections), and these points were used to select regions and decide if a crypt was present on that region (see Figures S1f and S2f).

#### (i) Parameterisation

We defined a set of main parameters that helped defining a crypt-like section automatically. These parameters were the number of harmonics required for the boundary approximation, the potential budding section area and arc length. However, for the last two parameters, we decided to consider normalised values according to the total organoid value, as these will vary according to the size of organoid image:

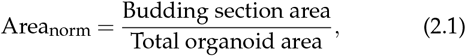

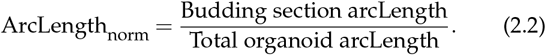

Our approach therefore accounts for the expected budding size range according to the experimental day. Each day should have its optimal value of the minimum and maximum normalised area (*Area*_*norm*_), allowing us to discriminate noise and villi regions from crypt regions. Additionally, a threshold for minimum normalised arc length (*ArcLength*_*norm*_) was included to avoid areas with sharp corners that are not present in crypts.

#### (ii) Training

Our experimental data consists of a total of 69 images: 30 of day 3, 19 of day 5 and 20 of day 7. The selection of best-fitted parameters was calculated by training the code using a selection of 5 images of each experimental day, which contained a different number of organoids (i.e. day 3 data contained 18 organoids) in the case of *in-vitro* data. In the case of *in-silico* data, a group of 10 organoids per simulated day was selected for training. All the selected organoids’ crypts were manually counted as a control during the training phase.

An objective function was generated for the training and parameter optimisation. This function included the code-calculated crypts, the hand-counted crypt values, and the calculation of percentage error comparing these two values:

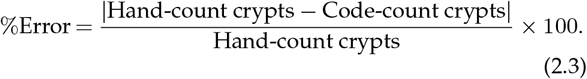

This objective function was used to find the global minima using the simulated annealing algorithm within MATLAB’s Global Optimisation Toolbox. This method was selected due to its capacity to explore globally for parameter values and prevent the system from being trapped in local minima in early iterations.

The optimised parameters obtained from the training are displayed in Table 2. In general, the training data suggest that the code can have around ∼80% of accuracy, so it can be used to approximate the mean crypt count at different time points of an intestinal organoid experiment.

**Table 2:** Optimised parameter values. Results obtained during optimisation of parameter values to identify crypts on both *in-vitro* and *in-silico* organoids

| Simulated day | Parameters |  |  |  | Error |
| --- | --- | --- | --- | --- | --- |
|  | Harmonics | Min crypt area | Max crypt area | Min arc length |  |
| In-vitro organoid images: OrganoSeg |  |  |  |  |  |
| Day 3 | 9 | 0.0638 | 0.2017 | 0.0290 | 14.3333 |
| Day 5 | 15 | 0.0259 | 0.1390 | 0.0630 | 17.9303 |
| Day 7 | 15 | 0.0176 | 0.1180 | 0.0828 | 20.7460 |
| In-vitro organoid images: Manual segmentation |  |  |  |  |  |
| Day 3 | 7.8 | 0.0666 | 0.2736 | 0.1466 | 16.8056 |
| Day 5 | 14.6 | 0.0153 | 0.2796 | 0.0784 | 22.1275 |
| Day 7 | 18.9 | 0.0121 | 0.5567 | 0.0737 | 22.5834 |
| In-silico organoid images: Both original and proposed models |  |  |  |  |  |
| Day 3 | 13 | 0.0079 | 0.0459 | 0.1000 | 17.5926 |
| Day 5 | 27 | 0.0052 | 0.0172 | 0.0576 | 15.7500 |
| Day 7 | 24 | 0.0068 | 0.0474 | 0.0357 | 12.8571 |

The optimised parameters were applied to calculate the number of crypts on the manually segmented organoids (n=132 organoids per day) and the organoids obtained with the OrganoSeg algorithm (n=124 for day 3, n=122 for day 5, n=109 for day 7). Additionally, as a control, we counted the crypts from the manually segmented organoids. For the *in-silico* data, on both the ‘original model’ and the ‘proposed model’, we had 50 simulated organoids per experimental day.

### (e) Circularity

Previous studies have used circularity as an indirect method to measure the growth of intestinal organoids [25, 30, 39–41]. Circularity has been regarded as an effective method due to the initial spheroid shape of organoids. And their subsequent production of crypts at later stages produces a loss of “roundness” or circularity. Therefore, it can be expected to obtain an almost perfect circle when the organoid has not yet produced crypts (circularity ≃ 1) and to lose this roundness once crypts appear (circularity *<* 1). Here, we will compare the results obtained from the presented crypt count method and the circularity obtained from the same images. Circularity can be calculated as:

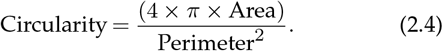

All p-values shown in Figures 2, 6 and S4 are between each indicated set of bars, and were calculated using a two-sample t-test.

**Figure 2:**
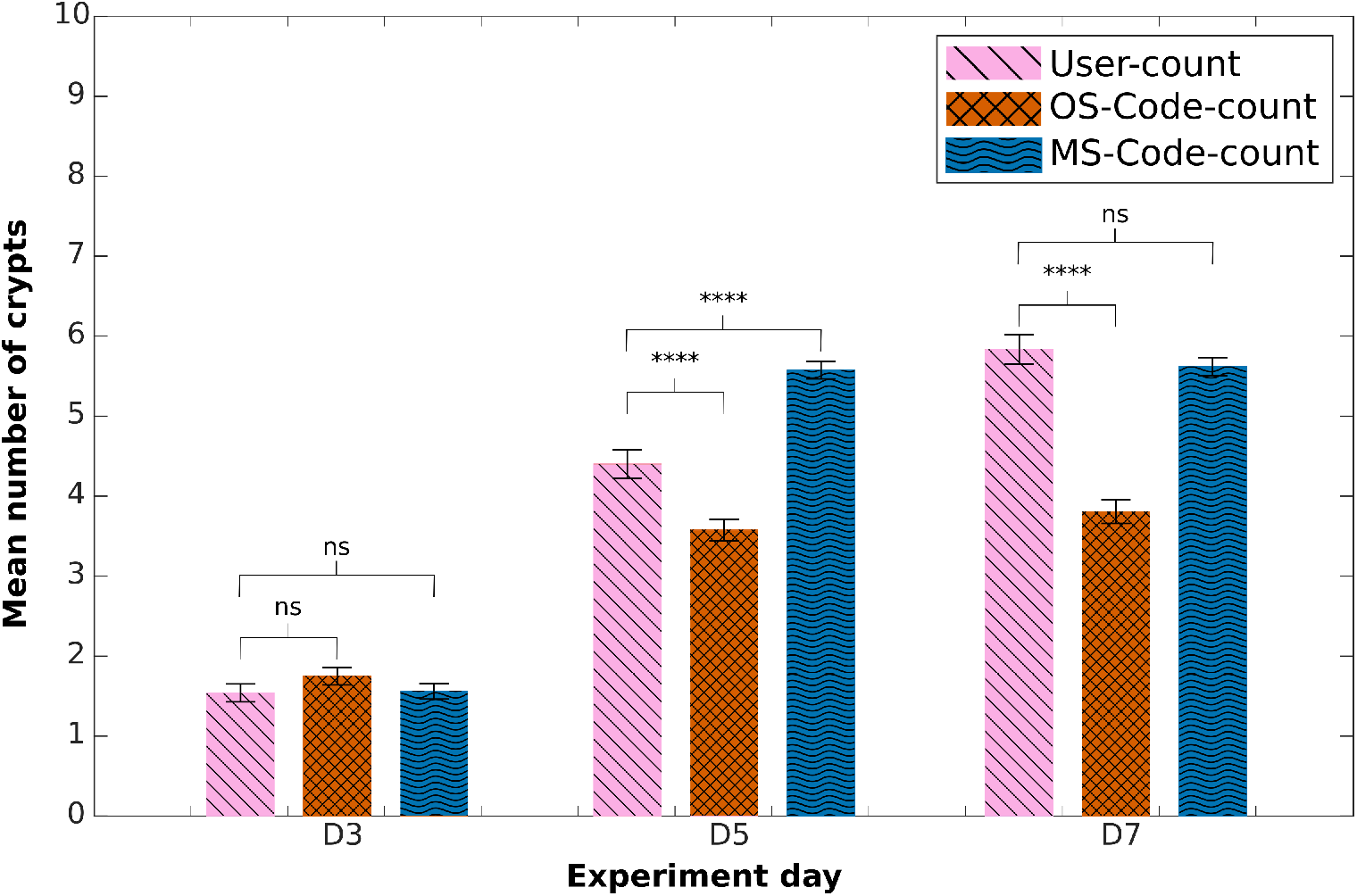
Mean number of crypts in experimental images. Comparison of mean number of crypts obtained using the counting-crypts code applied to the OrganoSeq- or manually-segmented images (‘OS-Code-count’ and ‘MS-Code-count’, respectively). The ‘user-counted’ refers to manually counted crypts. P-values from two-tailed unpaired t test computed over the data sets shown, ∗p<0.05, ∗∗p<0.01, ∗ ∗ ∗p<0.001, ∗ ∗ ∗∗p<0.0001

## 3. Results

Figure 2 exhibits the results of the mean number of crypts obtained from manual user count (‘user-counted’) and the code-counted using the previously presented segmentation methods, namely, OrganoSeg and Manual segmentation. To avoid confusion we will name the results obtained from applying the crypt-count algorithm to masks obtained from OrganoSeg and Manual segmentation ‘OS-Code-count’ and ‘MS-Code-count’ respectively. Here, it can be observed that the code-counted results obtained from ‘OS-Code-count’ and ‘MS-Code-count’ are not significantly different at day three compared to ‘user-counted’ data. The overall trend among the three data sets shows similar organoid growth. However, in data sets where the code-counting crypt algorithm was applied, we can observe a more significant increase in the mean number of crypts between days 3 and 5; and a minor growth change between days 5 and 7.

Results obtained from ‘MS-Code-count’ show no significant difference on days 3 and 7 compared to ‘user-counted’ data. Nonetheless, results from day 5 show a substantial difference between these two sets. Figure S3 displays the histograms and box plots collected for this analysis. Here it is shown that in experimental day 5 (S3b/S3e) we obtained less overlap of found crypts per organoids than the one observed for days 3 and 7, displayed in Figures S3a/S3d and S3c/S3f.

These results suggest that segmentation accuracy is essential to improve the performance of the code-counting crypt program. In the case of ‘MS-Code-count’, possible factors that limit the accuracy at day 5 include the variability in crypt sizes or the inability of the optimisation function to obtain better parameters at this stage. As mentioned in the methods section, the code is limited to a specific set and parameter range that may not be as flexible to cover the variability in organoid morphology present at this point of the experiment.

Next, we applied the crypt-counting algorithm to the *in-silico* organoids obtained using the Langlands et al. model. Results in Figure 3 display that the ‘original model’ is capable of replicating organoid growth in terms of crypt count at day 7. However, there was room for improvement to replicate the growth rate observed on days 3 and 5. Therefore, we decided to perform some modifications to the model to make it more biologically realistic and observe if these modifications were enough to improve the growth rate of the number of crypts in the *in-silico* organoids.

**Figure 3:**
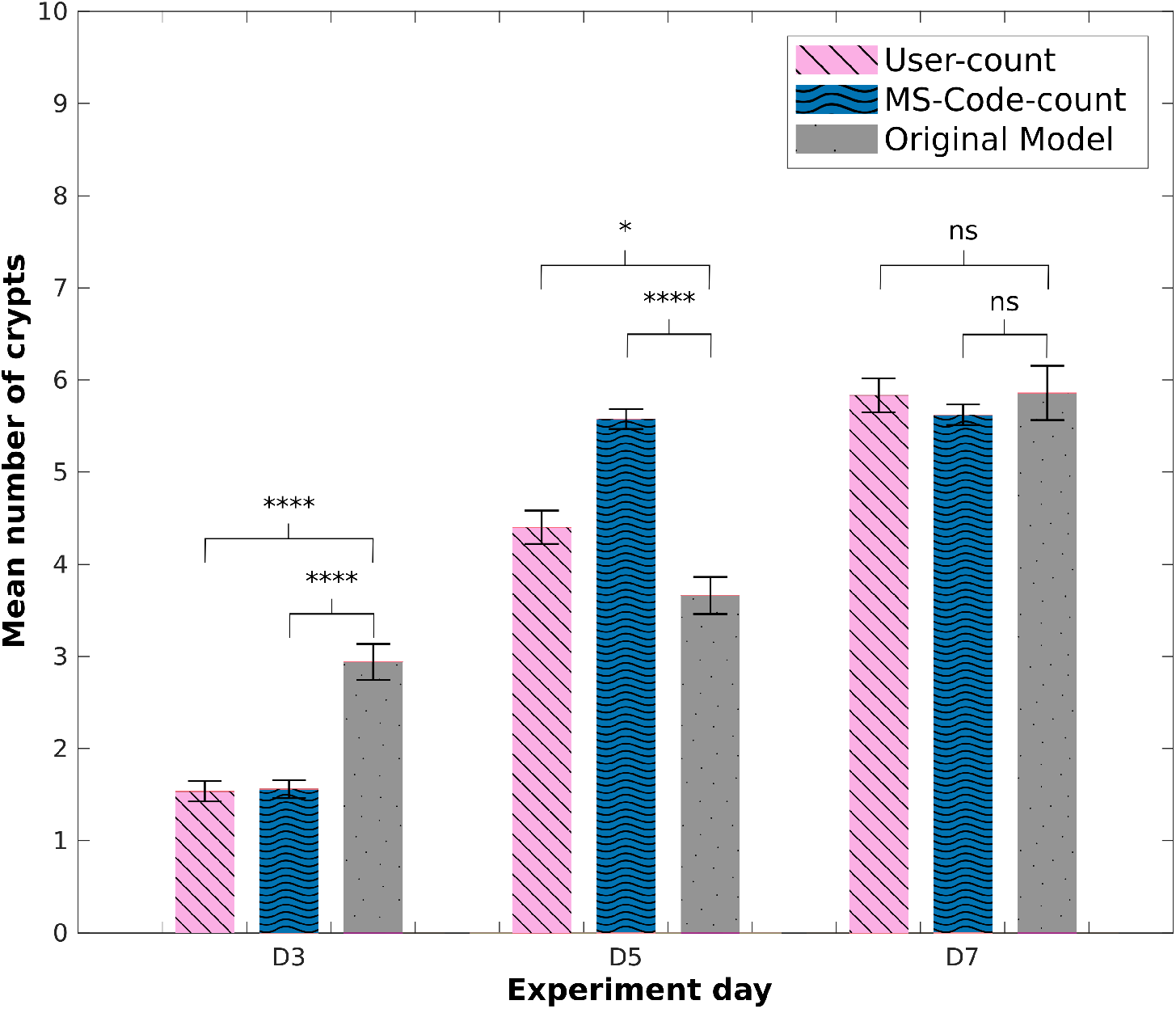
Mean number of crypts: Original *in-silico* model. Comparison of mean number of crypts obtained using the counting-crypts code applied to manually segmented *in-vitro* data (‘MS-Code-count’), *in-silico* organoids obtained from the ‘original model’ and the ‘user-counted’ crypts. P-values from two-tailed unpaired t test computed over the data sets shown, ∗p<0.05, ∗∗p<0.01, ∗ ∗ ∗p<0.001, ∗ ∗ ∗∗p<0.0001

### (a) Proposed modifications to the *in-silico* model

#### (i) Initial conditions

As previously mentioned, the *in-silico* organoids are compared to the control *in-vitro* experiment using crypts extracted from mouse intestine. Several studies mention that the intestinal organoids contain similar cell proportions that those found *in vivo* [4, 17, 42–45]. A study by Haber et al. [46] reported the profiling of 53,193 individual intestinal cells in the small intestine of 7- to 10-week-old mice. The authors classified and quantified the presence of several cell types as enteroendocrine (EEC), enterocyte (EnC), enterocyte progenitor (EnC-P), transit-amplifying cells (TA) (including on S and M cell-cycle phases), Paneth (PC), stem (SC), goblet (GC) and Tuft (TC) cells. Here, we extracted the cell population numbers of only the stem, TA and Paneth cells found in this study and inferred the initial proportion for the model. We assume that only those cell types are present in the system at the start of the simulation, as the experiments initiate from crypt fractions, and those cell populations mainly reside there. Thus, the initial cell population proportions used on all simulations of the ‘proposed model’ were: 42% stem cells, 49% TA cells and 9% Paneth cells. On the other hand, the initial proportions set for the simulations performed with the initial model were directly related to the target proportion set for cell production too, which in this case was ∼ 20 stem cells and ∼ 80 Paneth cells.

We decided to address the limitations we observed in the initial model and performed a few modifications to observe their effects on the resulting simulations. First, the initial number of cell-nodes in the system was increased from 20×20 to 36×40, to allow greater area for the organoid’s growth during more extensive simulations. Additionally, the initial organoid centre was moved to fit the centre of the new mesh.

#### (ii) Cell production model modifications

In the previous model, a hard cell could divide and produce a soft cell, and vice-versa. However, if we associate stiffness with a specific cell type, their division would be restricted to their lineage. As previously mentioned, stem (SCs) and Paneth cells (PCs) have been previously represented as soft and hard (i.e., stiff) cells, respectively. Nevertheless, Paneth cells are terminally differentiated and do not divide [47]. Suppose the cell production relies only on stem cells in the simulation with time. In that case, the system will be saturated by the stiffer cells that cannot divide, as soft cells will be expelled from the epithelial monolayer.

Therefore, we decided to introduce two new cell types. The first was defined as transit-amplifying cells (TAs) to represent the possibility of having a highly replicative cell type as an hard cell. A second cell type was defined as a general differentiated enteroid cell (ECs) that represents all other differentiated cells that TAs can produce. It is assumed that PCs, TAs and ECs are all stiff cells, with a 4.5 greater stiffness ratio than SCs.

According to this, the simulated cell cycle was modified to allow stem cells to symmetrically divide and generate other stem cells, which simulates the maintenance of multipotency; or to asymmetrically divide and produce a TA or Paneth cell daughter (see Figure 4b). The stem cell division will be stochastic, depending on a specific target proportion set. In this study, the probability proportions for the production of each cell type derived from stem cells were assumed to be:

**Figure 4:**
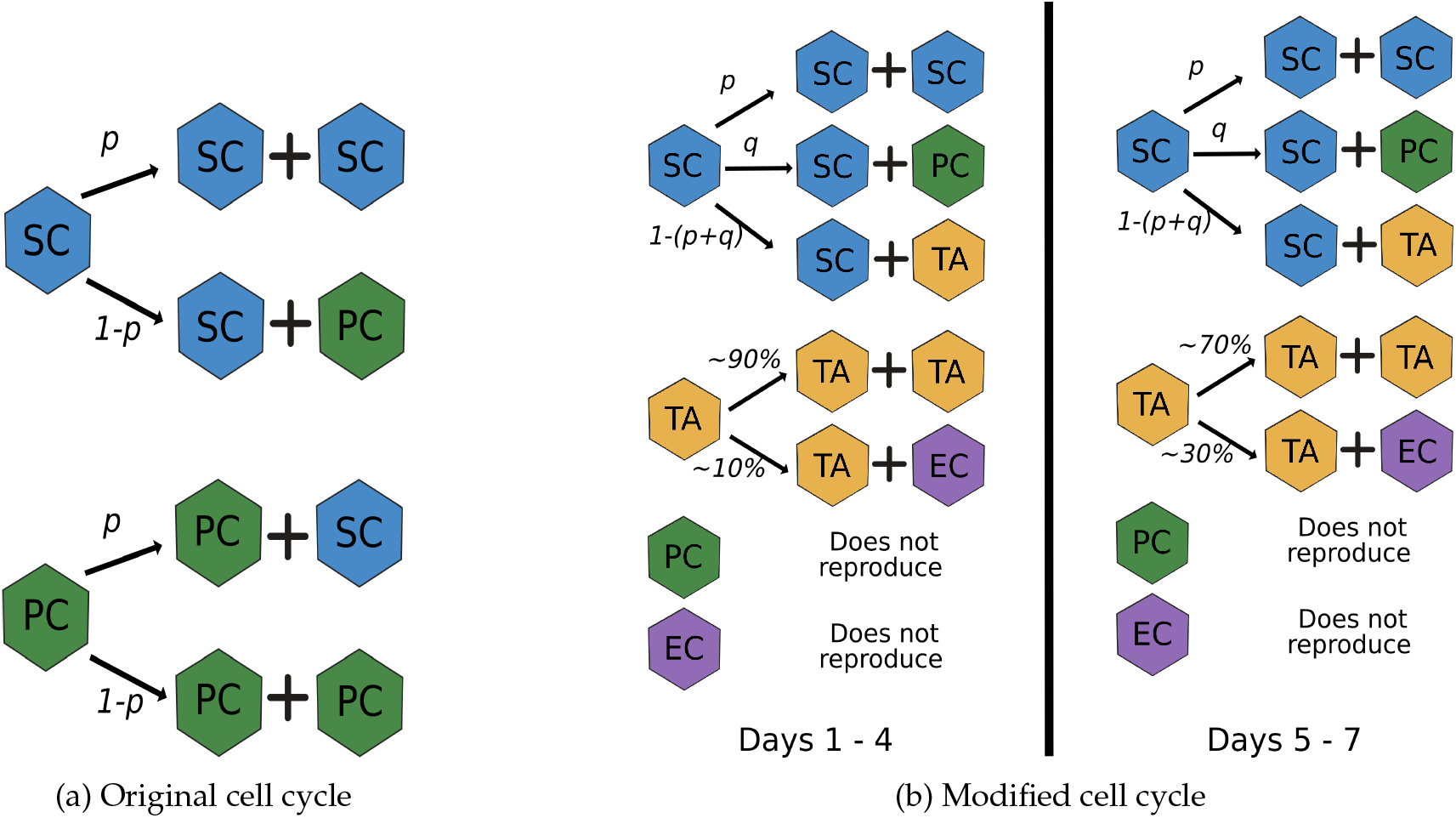
Cell cycle diagrams. (a) Original cell-cycle model used by Langlands et al. and Almet et al. [25, 30], (b) ‘proposed model’ modification that requires a production probability for each cell type derived from stem cells. Both models maintain the stochasticity of the system, as the selection of cell type is dependant on a random number generated at each time-step. Cell types: stem cell (SC); Paneth cell (PC); transit amplifying cell (TA); enterocyte (EC).

- ∼ 89% other stem cell;
- ∼ 9% a Paneth cell;
- ∼ 2% a TA cell daughter.

Using these proportions, we could regulate the cell production in the model. The mentioned proportions were selected as an educated guess, as PCs consist of around ∼ 9% of the cells in the intestine [46] and the model assumes that SCs are their only source. Furthermore, the probability of SC symmetric division (SC prop) was set to (∼ 89%) based on a study performed by Itzkovitz et al. [48], in which they observed a high symmetrical division of Lgr5+ cells on small crypts. On the other hand, the proportion of TAs obtained from SCs (TA prop) was set to be low (∼ 2%) to avoid overpopulation of this cell type, as TAs already in the system will produce more TAs by symmetrical division. On the other hand, TA cells can divide symmetrically, according to a higher probability proportion during the first five simulated days, as it has been reported that TAs undergo a limited series of cell divisions before differentiation [49, 50].

Thus, we assumed that during the first five simulated days of organoid growth, there is a higher probability of TAs producing more TA daughters. Then, the probability of TAs producing ECs is increased after this period. ECs, as well as PCs, are considered final differentiated cells and do not divide (see Figure 4b).

The probability proportions for TAs cell division are assumed to be:

- Before day 5
  – ∼ 90% other TA daughter
  – ∼ 10% ECs daughter
- After day 5
  – ∼ 70% other TA
  – ∼ 30% ECs daughter

A sweep of TAs and ECs probabilities was performed (not shown); the values presented in this study resulted in the most similar crypt count results compared to our *in-vitro* experiments. Figure 5 displays images of the simulated organoids using either the original (i.e. [25, 30]) and our ‘proposed model’.

**Figure 5:**
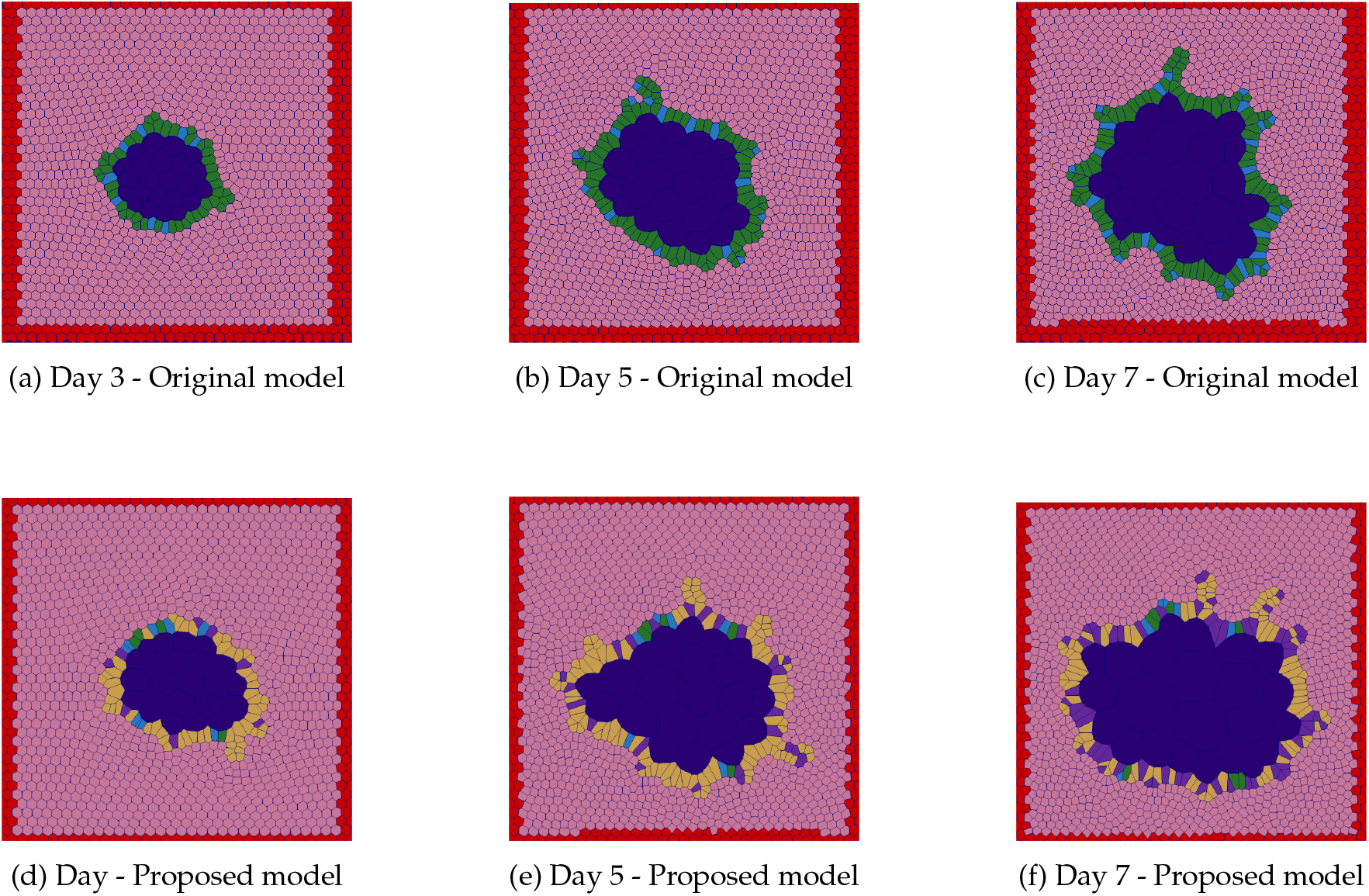
Timeline of *in-silico* organoids. Samples of simulation results at days 3 (a,d), 5 (b,e) and 7 (c,f). The ‘original model’ results are shown from a to c; and the ‘proposed model’ results are presented from d to f. Colour code: Stem cells (blue), transit amplifying cells (yellow), Paneth cells (green), Matrigel™ (pink), lumen (dark blue) and simulation boundary (red).

Results obtained from the mean number of crypts from the ‘user-counted’ show an exponential increment as the experiment progresses. This effect is displayed in the *in-silico* results obtained using the ‘proposed model’. Figure 6 presents a comparison of the ‘user-counted’, the ‘MS-Code-count’ and the code-counted crypts obtained from the ‘proposed model’. The results obtained from simulations on day 3 present a higher average number of crypts compared to the number of crypts found in the experiment, similar to what was observed previously in the ‘original model’. Nonetheless, for day 5, we observed that the ‘proposed model’ was able to replicate the mean number of crypts as the *in-vitro* data quantified by the user. Similarly to what was observed in the initial model, results obtained from day 7 present a similar average number of crypts per organoid across *in-vitro* sets.

**Figure 6:**
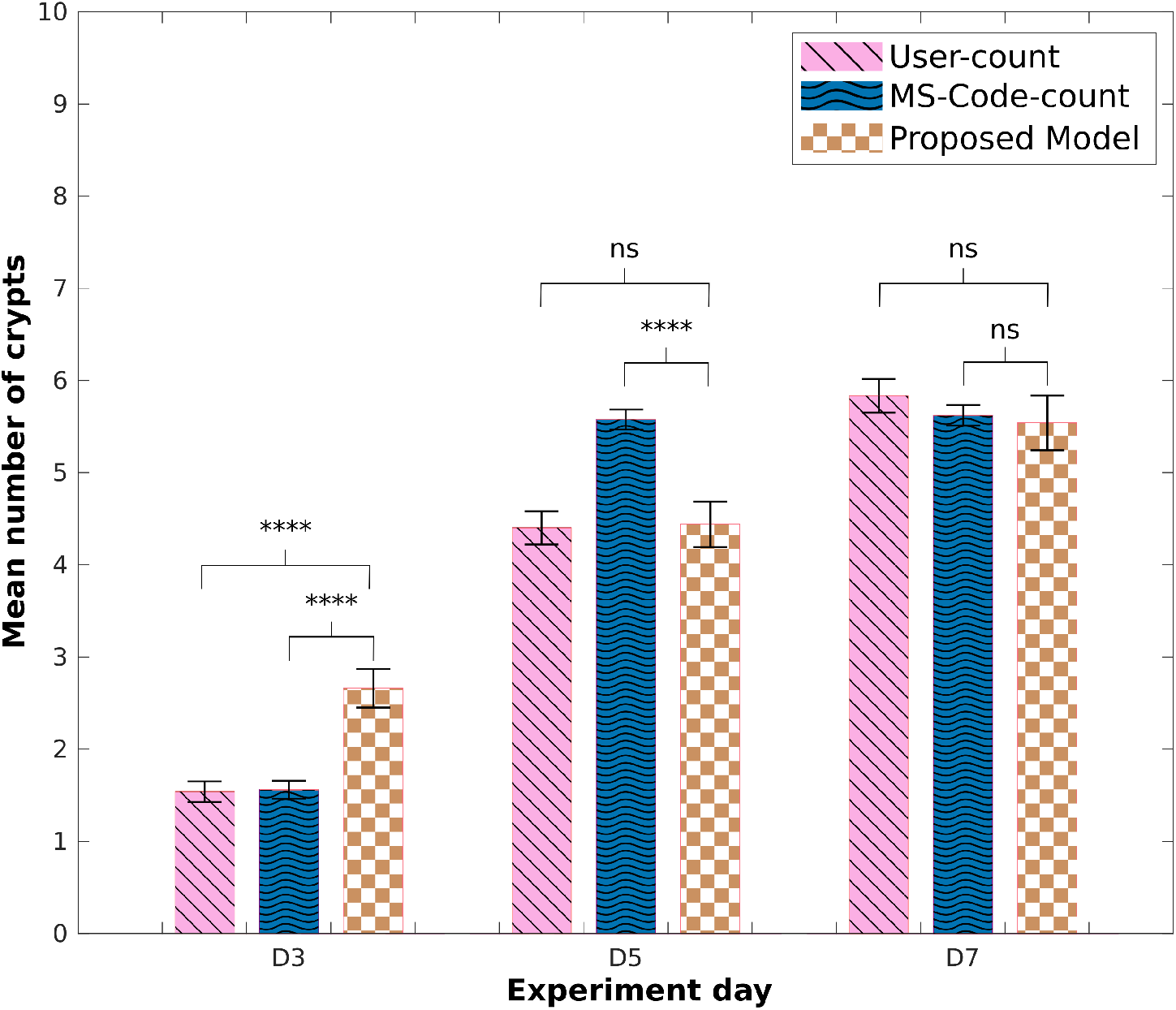
Mean number of crypts: Modified *in-silico* model. Comparison of mean number of crypts obtained using the counting-crypts code applied to the *in-silico* organoids simulated with the ‘proposed model’, and to manually segmented *in-vitro* data (‘MS-Code-count’); ‘user-counted’ crypt numbers are also shown. P-values from two-tailed unpaired t test computed over the data sets shown, ∗p<0.05, ∗∗p<0.01, ∗ ∗ ∗p<0.001, ∗ ∗ ∗∗p<0.0001

Finally, we were interested in observing how our crypt-count results compare to traditionally used methods. Figure S4 and Table 3 show the results obtained from measuring the circularity of organoids through manual segmentation on experimental data, and on simulated data (from the Original [25, 30] or our model). We should expect reduced circularity with an increased number of crypts as time progresses. Instead, the mean circularity obtained from manually segmented images of *in-vitro* organoids shows an increment, which suggests greater roundness, from day 5 to day 7. Similarly, the simulations obtained from the ‘proposed model’ do not show a difference in circularity as organoids grow.

**Table 3:** Mean circularity vs mean number of crypts. Results obtained from *in-vitro* and *in-silico* data obtained from both models. The circularity was calculated using equation 2.4, and here the manually segmented images were used. The mean number of crypts per organoid was obtained by applying the counting-crypts code to the resulting simulations of both models (original and proposed). The mean number of crypts calculated using the models is being compared to *in-vitro* user counted crypts. The standard error of the mean (SEM) was calculated for each set.

|  | Day 3 |  |  | Day 5 |  |  | Day 7 |  |  |
| --- | --- | --- | --- | --- | --- | --- | --- | --- | --- |
|  | User count | Original model | Proposed model | User count | Original model | Proposed model | User count | Original model | Proposed model |
| **Mean circularity** | 0.7813 | 0.7795 | 0.7597 | 0.5742 | 0.6278 | 0.5220 | 0.6487 | 0.5674 | 0.5217 |
| **SEM** | 0.0113 | 0.0084 | 0.0132 | 0.0101 | 0.0097 | 0.0139 | 0.0088 | 0.0106 | 0.0177 |
| **Mean no of crypts** | 1.5379 | 2.9400 | 2.7200 | 4.4015 | 3.6600 | 5.1000 | 5.8333 | 5.8600 | 6.4600 |
| **SEM** | 0.1120 | 0.1946 | 0.1853 | 0.1802 | 0.2012 | 0.2832 | 0.1829 | 0.2956 | 0.3335 |

## 4. Discussion

Cell and tissue imaging and image analysis can be instrumental to measure and quantify complex cell phenotypes in time and space. There has been a rise in the development of automatic and semi-automatic algorithms that specialise in the segmentation of two and three-dimensional cell cultures, including those based on label-free images [13, 51–54].

In the case of organoid cultures, algorithms have recently been proposed to segment and perform basic morphological image analysis by using traditional segmentation methods [32], or based on organoid staining [55]; other methods leverage machine-learning/deep learning approaches [56–59]. Nevertheless, these software are yet to be improved to detect and measure unique structures for specific organoid tissues. For example, in the case of intestinal organoids, the appearance and number of buds that replicate the shape and function of intestinal crypts.

Previous studies have counted crypts or buds on intestinal organoids by staining or labelling stem cells and segmenting regions containing a group of these cells to automatically or manually count them [60–64]. In cases where no labelling or staining is implemented during experimentation, we still rely on the user to define and count the number of crypts per organoid. Thus, we believe there is an opportunity for new software targeted at quantifying morphological structures on label-free bright-field images of organoids. This measure, especially on intestinal organoids, can be essential to define organoid cultures’ proper growth and standardisation.

The crypt-counting algorithm developed in this study presents a valuable tool to calculate an average of crypt-like or buds structures present in their *in-vitro* cultures. The algorithm has proved capable of approximating the average budding structures found in *in-vitro* intestinal organoid cultures on days three and seven after seeding. Nonetheless, the code shows limitations in calculating the number of crypt-like structures in sample dates in which there is a higher variability of organoid growth in terms of the number of budding sections. Also, in the case of the crypt-counting algorithm applied to *in-silico* images, we can observe a mismatch in the mean number of crypts compared to the *in-vitro* data. On day three, organoids start breaking symmetry from their original spheroid shape. Thus, it could be challenging to distinguish a crypt forming in the early stages, which can lead to an under- or over-counting of crypts, as the training depends on the user expertise. Additionally, the versatility of the algorithm to be applied to agent-based simulations of organoids provides the opportunity to improve such models and make predictions of pattern or structure formations before experimentation, which has been tested before for pattern formation in 2D stem-cell cultures [65]. In the future, the code limitations could be improved by implementing deep learning and machine learning algorithms to aid in finding additional or better-fitted parameters to define crypt-like structures [57, 66, 67].

It is important to notice that circularity was used before to test the accuracy of the ‘original model’. However, the results obtained in this study are not the same as those presented on [25]. Their circularity values calculated for 100 simulated hours are lower than those obtained in the present study. This may be due to the different methods used to extract and approximate the boundary from the simulations. Here we performed the same image extraction and boundary analysis to both *in-vitro* and *in-silico* images, in contrast to calculating the simulated epithelial layer circularity directly from the model’s cell connections. Additionally, the ‘original model’ did not present issues of cell competence or the removal of soft cells from the simulation due to their cell production model, in which any cell can produce a daughter of any other cell type according to the target proportion set by the user.

The proposed changes to the *in-silico* model maintain the potential to produce simulations that replicate the number of budding structures found on *in-vitro* experimentation. Nevertheless, the model still presents limitations, such as the lack of signalling pathways that regulate cell production [68], and it is limited to the 2D structure of the epithelial layer. The spatial restriction affects the position of new cells added to the system during the simulations. Thus, we were not able to recreate an accurate location of the cell-type populations found on smell-intestine crypts, in which SCs and PCs are positioned at the bottom. Similarly, the framework structure gives rise to the expulsion of softer cells more easily than hard cells, which may prompt the model to end with fewer soft cells (i.e. Stem cells) at the end of the simulation.

## 5. Conclusions

The present study is limited to snapshots at three different days, with no tracking of organoids, which makes it difficult to observe their growth directly. Nevertheless, the present crypt-count code and *in-silico* model provide valuable tools with which to explore the effect of crypt development on intestinal organoids and stepping stones for future approaches to classifying morphological structure in organoids. We do foresee an increasing adoption of agent-based models also in engineering biology, with the aim at designing novel 3D cellular structures and tissues, and possibly integrating a description of biomechanical interactions with detailed subcellular whole-cell processes [69].

## Data Accessibility

The scripts developed for this paper are available in the following repository: https://github.com/slmontes/SimpleCryptCount_Project.git

## Authors’ Contributions

S.M-O., M.H. and L.M. designed this research; S.M-O. implemented the mathematical models, generated simulations, analysed results and prepared the figures; S.M-O. and A.L. developed the algorithm; D.L. and A.C.W. performed the experiments and obtained the experimental images; A.G.F. supported the agent-based simulations; S.M-O., A.C.W., A.G.F., M.H. and L.M. wrote the manuscript; M.H. and L.M. supervised the project. All authors read and approved the final manuscript.

## Acknowledgements

The authors gratefully acknowledge the expertise provided by Dr Dominic Alibhai and Mr Adam Chambers from the Wolfson Bioimaging Facility, University of Bristol in the development of the initial custom code for image analysis. The authors also thank Dr Thomas Gorochowski and Dr Christopher Parker for their insightful comments and discussions throughout this work.

