## Supplemental material for "*In-silico* and *in-vitro* morphometric analysis of intestinal organoids"

### Supplementary Material

The following supporting material is composed of figures explaining the crypt counting algorithm functionality, and an additional presentation of the results obtained. The crypt counting algorithm and modified 2D model code are available online: `https://github.com/slmontes/SimpleCryptCount_Project.git`.

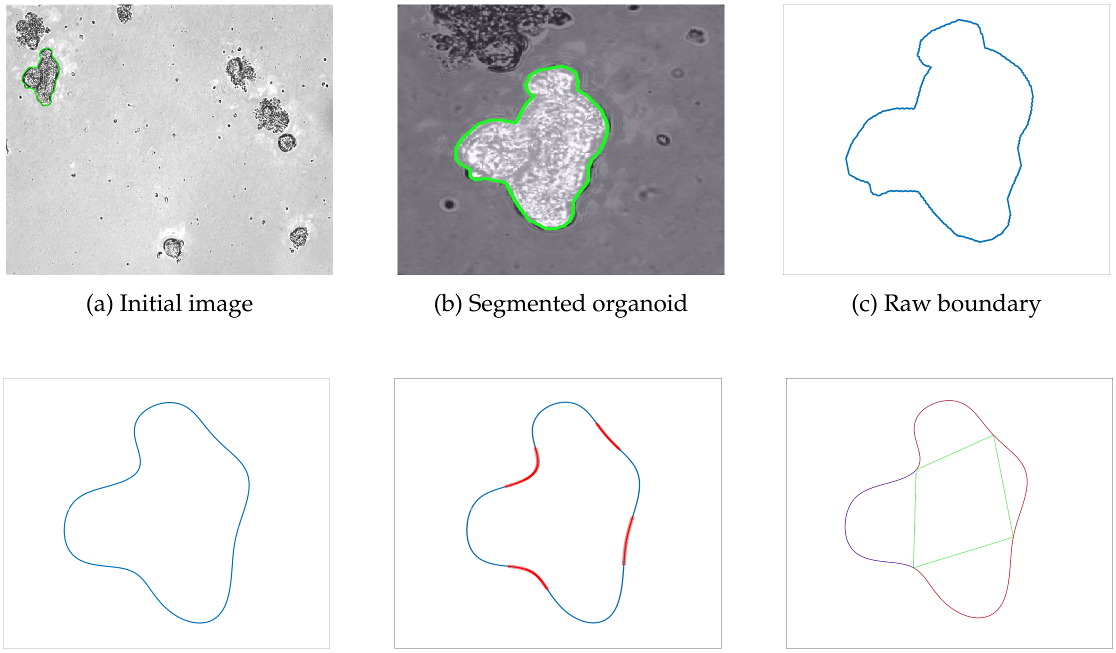

(a) Initial image (b) Segmented organoid (c) Raw boundary

(d) Approximated boundary (e) Inward boundary's sections (f) Tentative crypt sections

Figure S1: **Process performed on *in-vitro* images.** This image shows the general processing performed for each organoid's boundary to calculate the number of crypts present after extracting the mask's information. (a) Initial stacked image of a 3 day old organoid culture; (b) manual segmentation of an organoid; (c) raw boundary extracted from the segmented organoid; (d) boundary approximation obtained using a Fourier approximation; (e) calculated concave sections (in red) on the boundary; and (f) possible crypt sections detected by our algorithm.

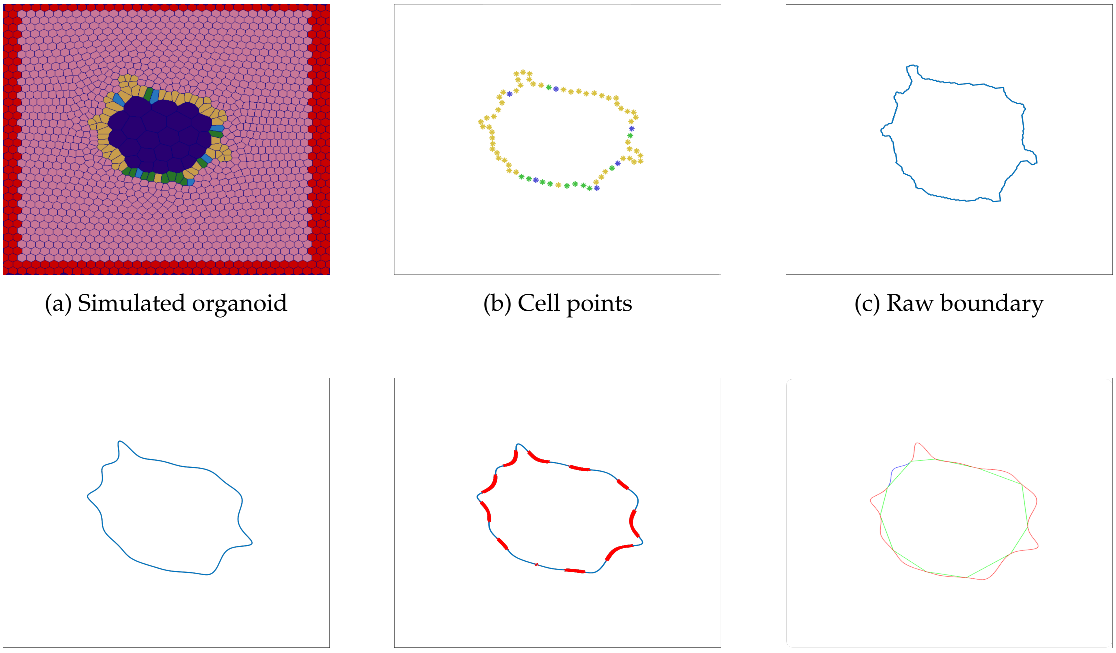

(a) Simulated organoid (b) Cell points (c) Raw boundary

(d) Approximated boundary (e) Inward boundary's sections (f) Tentative crypt sections

Figure S2: **Process performed on *in-silico* images.** This image shows the general processing performed for each *in-silico* organoid's boundary to calculate the number of crypts presented after extracting the mask's information. (a) Initial image obtained from a 3 day old organoid simulation with several nodes representing stem cells (blue), transit amplifying cells (yellow), Paneth cells (green), Matrigel™ (pink), lumen (dark blue) and the simulation boundary (red); (b) location of epithelial cells ( SC (blue), TA (yellow), PC (green) ) extracted from the simulation; (c) raw boundary of the simulated organoid; (d) boundary approximation obtained using a Fourier approximation; (e) calculated concave sections (in red) on the boundary; and (f) possible crypt sections detected by our algorithm.

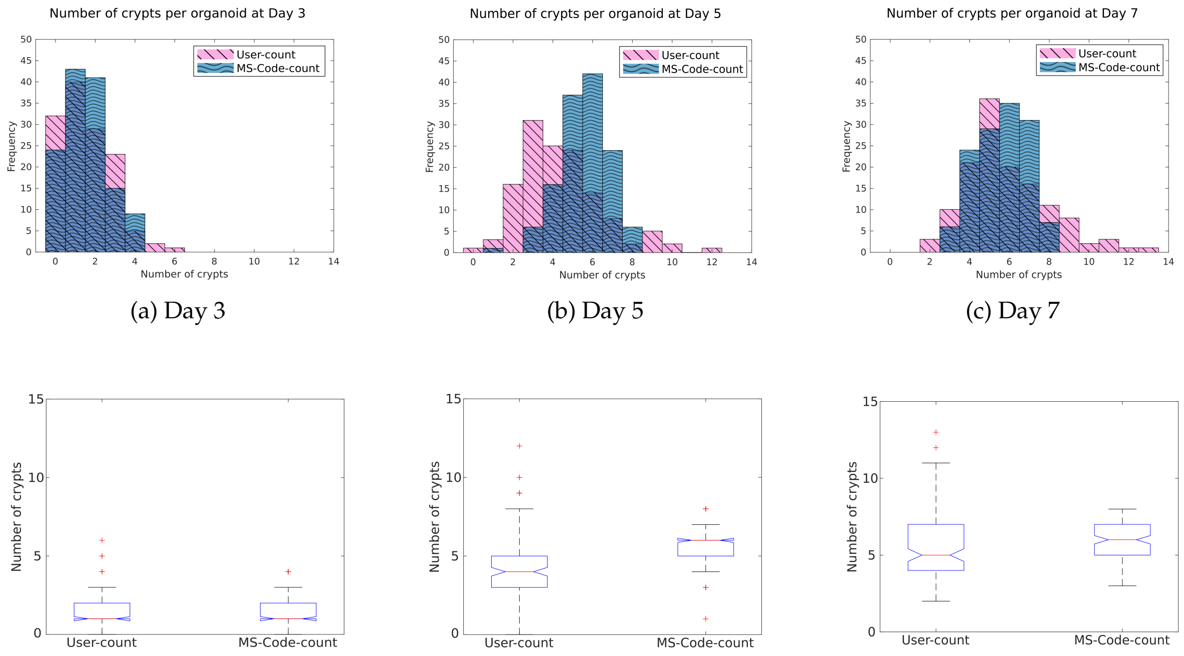


(a) Day 3 (b) Day 5 (c) Day 7

(d) Day 3 (e) Day 5 (f) Day 7

Figure S3: **Mean number of crypts found from *in-vitro* images: ‘user-counted’ vs ’MS-Code-count‘.** Comparison of number of crypts found per organoid at each day by the user in comparison to those found in ’MS-Code-count‘. (a-c) Histograms of the distribution of crypts found by the user (pink), compared to ’MS-Code-count‘ (blue). (d-f) Boxplots of the number of crypts found using the previously mentioned methods, in which the boundaries of the box represent the $25^{th}$ and $75^{th}$ percentiles respectively of the median (red line) number of crypts found per day, the width of the notch represents a 95% confidence interval around the median, the whiskers extend the most extreme data points, and the outliers are represented individually (red plus signs).

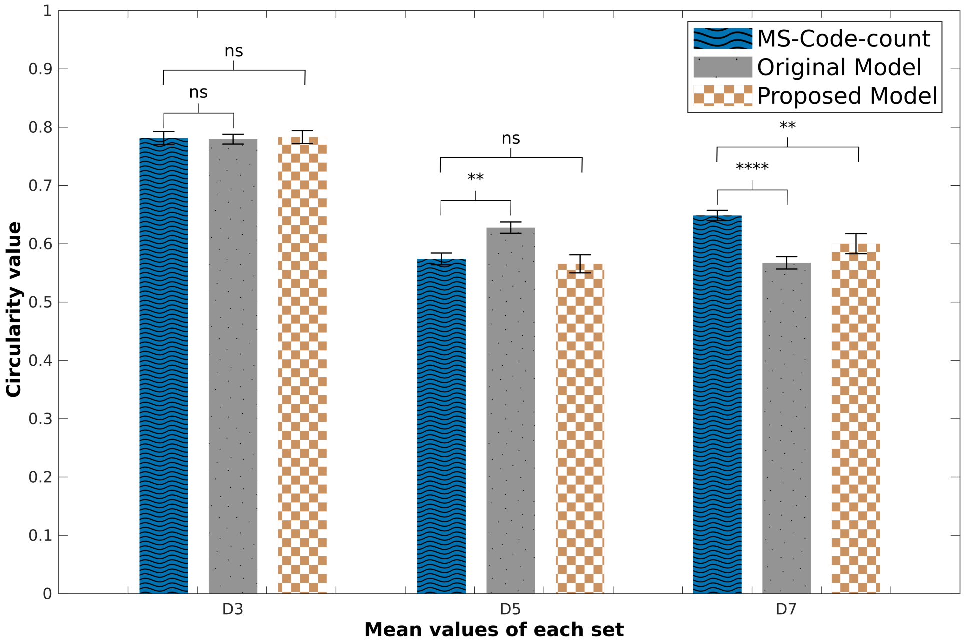


Figure S4: **Mean circularity.** Comparison of mean circularity calculated at each day for *in-vitro* (using the manually segmented images) and *in-silico* organoids. P-values from two-tailed unpaired t test computed over the data sets shown, $*p<0.05$, $**p<0.01$, $***p<0.001$, $****p<0.0001$
